# Upper Thermal Tolerance of the Endemic Freshwater Stingray *Potamotrygon Magdalenae* and Inference on its Populations: An Endemic Potamotrygonid Resistant to High Temperatures Facing Thermal Changes

**DOI:** 10.1101/2023.07.30.551136

**Authors:** Daniela Gómez-Martínez, Edgardo Londoño-Cruz, Paola Andrea Mejía-Falla

## Abstract

Knowledge on thermal tolerance limits provide important clues to the capacity of a species to withstand acute thermal conditions. Climatic models predict the increase and intensification of events such as heat waves, therefore understanding the upper thermal limits that a species can tolerate, has become of utmost importance. We measure the upper thermal tolerance of the endemic freshwater stingray ***Potamotrygon magdalenae*** acclimated to experimental conditions, using the Critical Thermal Methodology (CTM). We also describe the behavior of individuals and infer the possible consequences of temperature increases in the habitats of *P. magdalenae* populations. There were no significant differences between sexes in temperature tolerance or behavior. The Critical Thermal Maximum - CTMax (39ºC) was 5.9ºC above the maximum recorded temperature for the study area. Although *P. magdalenae* was tolerant to high temperature and currently is not living at its upper thermal limit, its survival in Guarinocito Pond will be threatened if the increasing trend in temperature conditions keeps growing over time.

**SUMMARY STATEMENT:** *Potamotrygon magdalenae* shows tolerance to high temperatures regardless of sex and size; however, can be threatened in the future if the temperature in its habitat continues rising.

## INTRODUCTION

Temperature is considered the abiotic master factor because of its profound effects on nearly every physiological process, including the limit to biochemical rates and geographic distributions (Di Santo and Bennett, 2011; Pérez et al., 2003). The level of those effects depends on thermal tolerance range of the species, defined as the “tolerance zone” or upper and lower temperature limits within which each species can survive or in which vital processes are not affected by thermal changes (Alfaro et al., 2005). These thermal tolerance limits can be calculated using the Critical Thermal Maximum (CTMax) and the Critical Thermal Minimum (CTMin), by exposing the individuals to a constant rate of increasing or decreasing temperature, until a sublethal or non-lethal point is reached, *i.e*., the level at which individuals start to display tetanic contractions of the pectoral disc, muscular spasms and unorganized locomotion (Beitinger et al., 2000, Dabruzzi et al., 2012). Thereby, the tolerance zone of the species delimits their ability to function under different environmental conditions, allowing to infer, among others, the possible effects of changes in habitat temperature on the populations characteristics. Such changes may be due to various phenomena, including El Niño or La Niña.

Fish temperature sensitivity and tolerance has been documented for several species (*e.g*. Lutterschmidt and Hutchison, 1997; Beitinger et al., 2000; Fangue and Bennett, 2003) and since its development in 1944 (Cowles and Bogert, 1944), the use of CTM to infer acute upper thermal tolerance in fishes continue to increase (Desforges et al., 2023). Despite these, in Colombia only four studies have been carried out on this subject (i.e. fish thermal tolerance), three of them on reef fishes in Gorgona National Natural Park, off the Pacific coast (Mora and Ospina, 2001; Mora and Ospina, 2002; Mora and Ospina, 2004) and one on *Poecilia caucana*, a fresh water fish (Martinez et al., 2016.)

On the other hand, several studies have shown that temperature affects the distribution of elasmobranchs populations (Almeida et al., 2009; Di Santo, 2015; Di santo, 2016). For instance, *Myliobatis californica, Triakis semifasciata* and *Mustelus henlei*, were all observed to shuttle among temperatures in Tomales Bay, California (Hopkins and Cech, 2003). Juvenile *Carcharhinus leucas* either permanently left an estuarine system or died during extreme “cold snap” in the Shark River estuary of Everglades National Park, USA (Matich and Heithaus; 2012). There are only four published studies on thermal tolerance of batoid fishes (Britton, 1924; Huntsman and Sparks, 1924; Fangue and Bennett, 2003; Dabruzzi et al., 2012), and none have been conducted on freshwater stingrays (Family Potamotrygonidae).

The freshwater stingray *Potamotrygon magdalenae* is found in the Magdalena basin and is the only endemic freshwater stingray in Colombia (Mejía-Falla et al., 2009). This species has ornamental use in some regions and high commercial value, being the most exported ornamental batoid from Colombia (60-70% of exported rays; Mejía-Falla et al., 2009; Lasso et al., 2013). It is nationally categorized as Near Threatened (Lasso et al., 2012) and internationally as Least concern (https://www.iucnredlist.org/species/161385/61472512, accessed 18.06. 23). It is also considered a Very High Priority research species in the National Plan of Action for the conservation and management of sharks, skates, rays and chimaeras of Colombia (Caldas et al., 2010). Investigation on this species includes studies on nephron structure (Ogawa and Hirano, 1982), morphology of intestinal epithelial cells (Teshima and Hara, 1983), enzymatic activity (Singer and Ballantyne, 1989), reproductive biology (Teshima and Takeshita 1992; Ramos-Socha and Grijalba-Bendeck, 2011; Pedreros-Sierra, 2015; Pedreros-Sierra et al., 2016; Anaya-López and Ramirez, 2017; Lizcano-Gutierréz and Ramírez – Pinilla, 2022), diet (Ramos-Socha and Grijalba-Bendeck, 2011), Feeding habits and ecological role (Márquez-Velásquez et al., 2019), Distribution and abundance (Castañeda et al., 2021), ecological and fisheries issues and aspects related to transport effects and reproduction and growth in captivity (Mejía Falla et al., 2016), Hematology and blood biochemistry under conditions of captivity (Pérez-Rojas J.G. et al., 2020). However, no studies related to thermal biology of this species have been carried out. These sort of studies are crucial to understand the thermal ecology of fishes, allowing the prediction of possible responses of these organisms to changes in their habitats (Dabruzzi et al., 2012). This study aimed to evaluate the maximum thermal tolerance of males and females of *P. magdalenae* and describe the behavioral response to thermal stress in order to answer if the populations of the Magdalena freshwater stingray in Guarinocito Pond is at the upper limits of its tolerance to high temperatures and infer the possible consequences of an increase in habitat temperature on its population.

## MATERIALS AND METHODS

### Capture and holding of individuals

The procedures and protocols used were approved by the Faculty of Natural and Exact Sciences from Universidad del Valle. Twenty-four animals (12 males and 12 females) were captured using cast-nets by local fishermen in Guarinocito Pond, La Dorada, Caldas (5º21’N, 74º44’W). Before beginning the experiment, stingrays were kept in floating plastic crates in the pond during 24 h; this allowed the animals to get used to confinement at temperatures similar to those of the water of the pond (24,5 – 33,1)°C. During the time of acclimation, the stingrays were fasted to avoid interference of feeding processes in responses to temperature changes (Coutant, 1977). After this, groups of three animals were selected randomly (males and females separately) to conduct the experiments (see next section). Each individual was allocated to either one control aquarium or to two temperature treatment aquaria. These aquaria were half filled with 30 l of freshwater taken directly from the pond and fitted with air pumps to allow continuous oxygenation and mixing of water.

### Experiments: Critical Thermal Maximum (CTMax) and behavior of individuals

In the treatment aquaria, temperature was gradually increased at a constant rate of 1°C/2 h with heaters electronically controlled, until a sub-lethal temperature was reached (*i.e*., the value at which individuals start to display tetanic contractions of the pectoral disc and a negative response to touch stimuli, equivalent to a non-response to the stimulus) (Beitinger et al., 2000; Dabruzzi et al., 2012). In the control aquarium, temperature was kept relatively constant and similar to the average temperature at the sampling site (mean ± SD = 31 ± 0,77°C). Since each experiment (i.e. 3 stingrays) was carried out independently and we only counted with three aquaria, it took eight days (24 stingrays* 3 stingrays ×day^-1^) to complete the survey. In total, 16 stingrays (8 males and 8 females) were subjected to temperature treatment and 8 were used as control (4 males and 4 females). It is important to clarify that in the control all data (observations) taken during the experiment (every two hours at the same temperature, 31°C) can be observed, while in the treatment the data were taken once per individual at each temperature.

Water parameters (temperature, conductivity, dissolved oxygen and pH) were monitored every two hours with a multiparameter probe (YSI Professional Plus) and a pH-meter (model HI 98127). Animal behavior was described every two hours (i.e. every 1°C increment) under both, temperature treatment and control conditions. Behavior was categorized as follows: lifting of the disc (null, low, intermediate, high; Fig. 1), degree of activity (Inactive: no movement was observed; Light activity: there were less than five movements in one direction; Active: there were five or less than five horizontal and vertical movements; Very active: there were over five horizontal and vertical movements accompanied by excitation), and response to first stimulus – touch – (None: when the animal did not respond to the stimulus; Slow: when the animal did not respond immediately to the stimulus; and Fast: when the animal responded immediately to the stimulus).

**Fig. 1.**
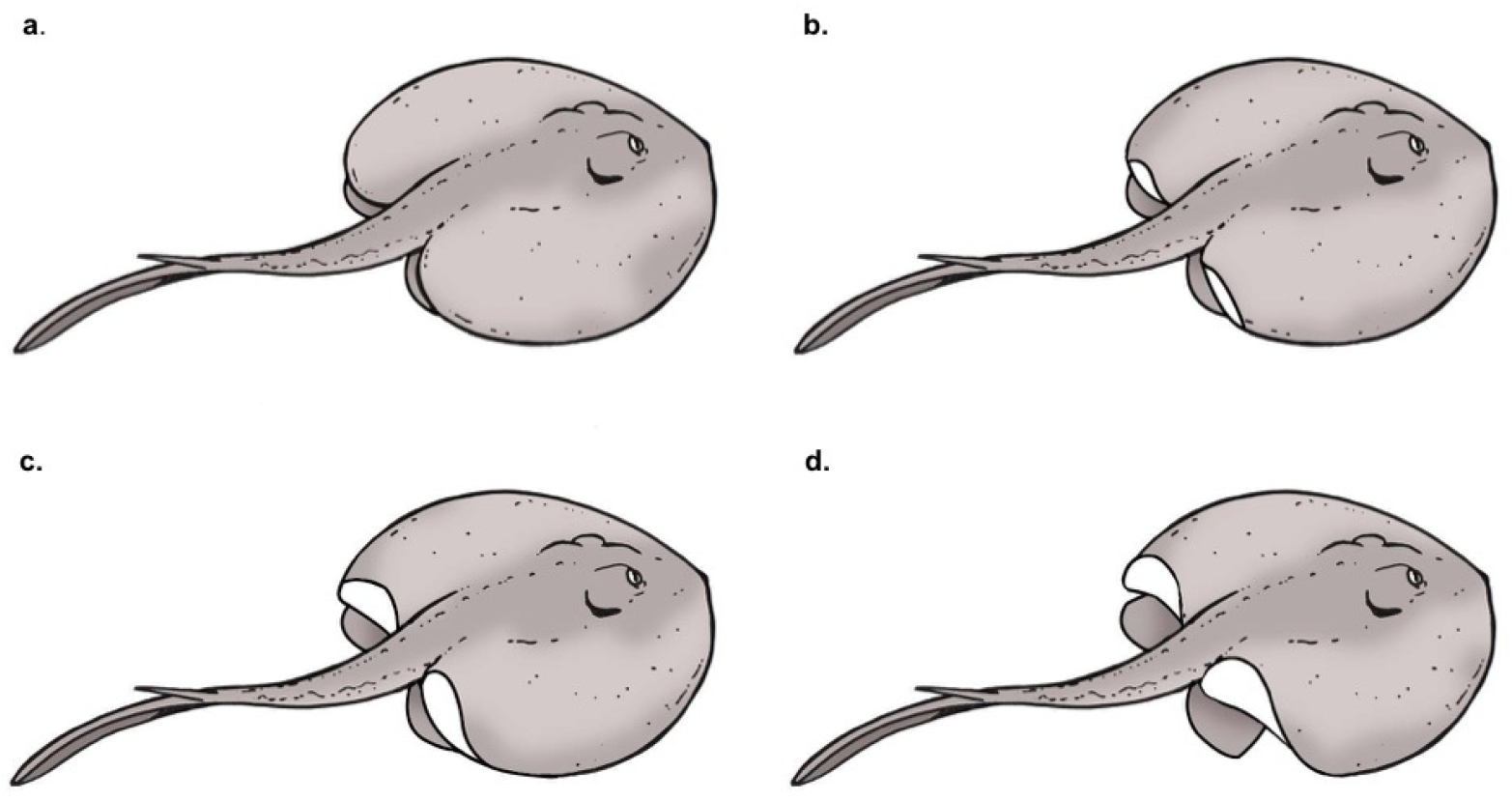
Representation of lifting of the disc of *Potamotrygon magdalenae* individuals in response to temperature increase. (A) Null, (B) Low, (C) Intermediate, (D) High.

Stingrays were measured (total length and disc width) after each trial to avoid stress from manipulation and returned to acclimation conditions. Once returned to the acclimation crates, stingrays were continuously observed to monitor their response to stimuli; once they were in normal condition, they were released.

### Data analysis

CTMax was calculated for all 16 individuals and for each sex as the arithmetic mean of the sublethal thermal points obtained in the treatment aquaria (Lowe and Vance, 1955; Beitinger et al., 2000). Differences of CTMax between the sexes were evaluated with a t-test, after normality and homoscedasticity was proved. Since there is slower heat penetration in larger animals (Hutchinson, 1976), we evaluated the relationship between animal size and CTMax with a linear regression; this test was not evaluated for each sex separately because no differences between CTMax between sexes were found. Differences in the parameters (conductivity, dissolved oxygen and pH) between control and treatment were evaluated with a Mann-Whitney test, to determine if any of these parameters can affect the response against a temperature increase. We also describe and discuss behavioral responses to acute thermal stress during the Critical Thermal Methodology (CTM).

### Relationship between CTMax of *Potamotrygon magdalenae* with local and regional temperatures

To infer whether the population of the Magdalena freshwater stingray in Guarinocito Pond is at the upper limits of its tolerance to high temperatures, we compared the obtained CTMax in treatment aquaria with maximum temperatures recorded in this pond and in others areas of the Magdalena River basin, where the presence of this species has been confirmed (data from SQUALUS Foundation and the Institute of Hydrology, meteorology and environmental studies, IDEAM; 2005 to 2014).

## RESULTS

Stingrays measured between 9,5 and 14,7 cm of disc width (DW) and between 20,3 and 35,3 cm of total length. The survival of individuals during the experiment was 100%.

### Critical Thermal Maximum (CTMax) of *Potamotrygon magdalenae*

The average (±SD) CTMax for the 16 specimens collected in Guarinocito Pond was 39.0°C (SD=±0,65). Although CTMax was higher in males (39,2±0,60°C) than females (38,8±0,67°C), there were no statistical significant differences (t=1,32, p=0,20). Moreover, CTMax and disc width were not correlated, neither for both sexes combined (r=-0,11, p= 0,65) nor for each sex separately (females: r=-0,14, p=0,71; males: r= 0,12, p=0,74).

The water parameters conductivity, dissolved oxygen and pH (Table 1) were similar between temperature treatment and control aquaria except for dissolved oxygen concentrations (n_c_=73, n_t_=145, U=1008,5, p=0,019).

**Table 1.** Ranges and averages of water parameters monitored.

| Treatment | Min.<br>pH | Max.<br>pH | Average<br>pH | Min.<br>DO<br>(mg/L) | Max.<br>DO<br>(mg/L) | Average<br>DO<br>(mg/L) | Max.<br>conductivity | Min.<br>conductivity | Average<br>Conductivity |
| --- | --- | --- | --- | --- | --- | --- | --- | --- | --- |
| Control<br>(31°C) | 7,48 | 8,56 | 8,26 | 1,0 | 2,5 | 1,64 | 232,0 | 105,3 | 199,02 |
| 31°C | 7,51 | 8,12 | 7,81 | 1,1 | 1,8 | 1,45 | 204,2 | 111,9 | 186,36 |
| 32 °C | 7,9 | 8,28 | 8,13 | 1,0 | 1,9 | 1,26 | 213,6 | 115,3 | 193,16 |
| 33 °C | 8,16 | 8,37 | 8,27 | 1,0 | 1,5 | 1,16 | 218,8 | 192,0 | 205,21 |
| 34 °C | 8,23 | 8,46 | 8,36 | 1,0 | 1,5 | 1,16 | 223,0 | 202,5 | 211,66 |
| 35 °C | 8,26 | 8,54 | 8,38 | 0,9 | 1,7 | 1,21 | 228,5 | 190,8 | 213,79 |
| 36 °C | 8,18 | 8,52 | 8,38 | 0,8 | 1,2 | 1,04 | 239,4 | 207,1 | 221,81 |
| 37 °C | 8,25 | 8,52 | 8,39 | 0,8 | 1,0 | 0,94 | 257,1 | 223,7 | 232,08 |
| 38 °C | 8,23 | 8,49 | 8,36 | 0,7 | 1,2 | 0,84 | 254,9 | 222,0 | 236,66 |
| 39 °C | 8,25 | 8,42 | 8,35 | 0,8 | 1,0 | 0,84 | 260,9 | 232,7 | 244,70 |
| 40 °C | 8,52 | 8,52 | 8,52 | 1,1 | 1,1 | 1,10 | 254,2 | 254,2 | 254,20 |

### Behavior of *Potamotryon magdalenae* individuals against the temperature increase

During the experiment, the lifting of the disc was observed in a 45,1% of the observations, with a higher percentage of occurrence in stingrays of the treatment (73,2%) than those in control (26,8%), and mainly (>60%) when the higher temperatures (>36°C) occurred. The low lifting of the disc was reduced (<30%) in both sexes, with the exception of most males at 36ºC (Fig. 2A,B) where 62,5% raised the disc at a low level. The high lifting of the disc (considered in this study as one of the two sublethal points) occurred exclusively in treatment aquaria and at temperatures above 38ºC in males (Fig. 2A) and above 35ºC in females, with the highest value at 39ºC (Fig. 2B).

**Fig. 2.**
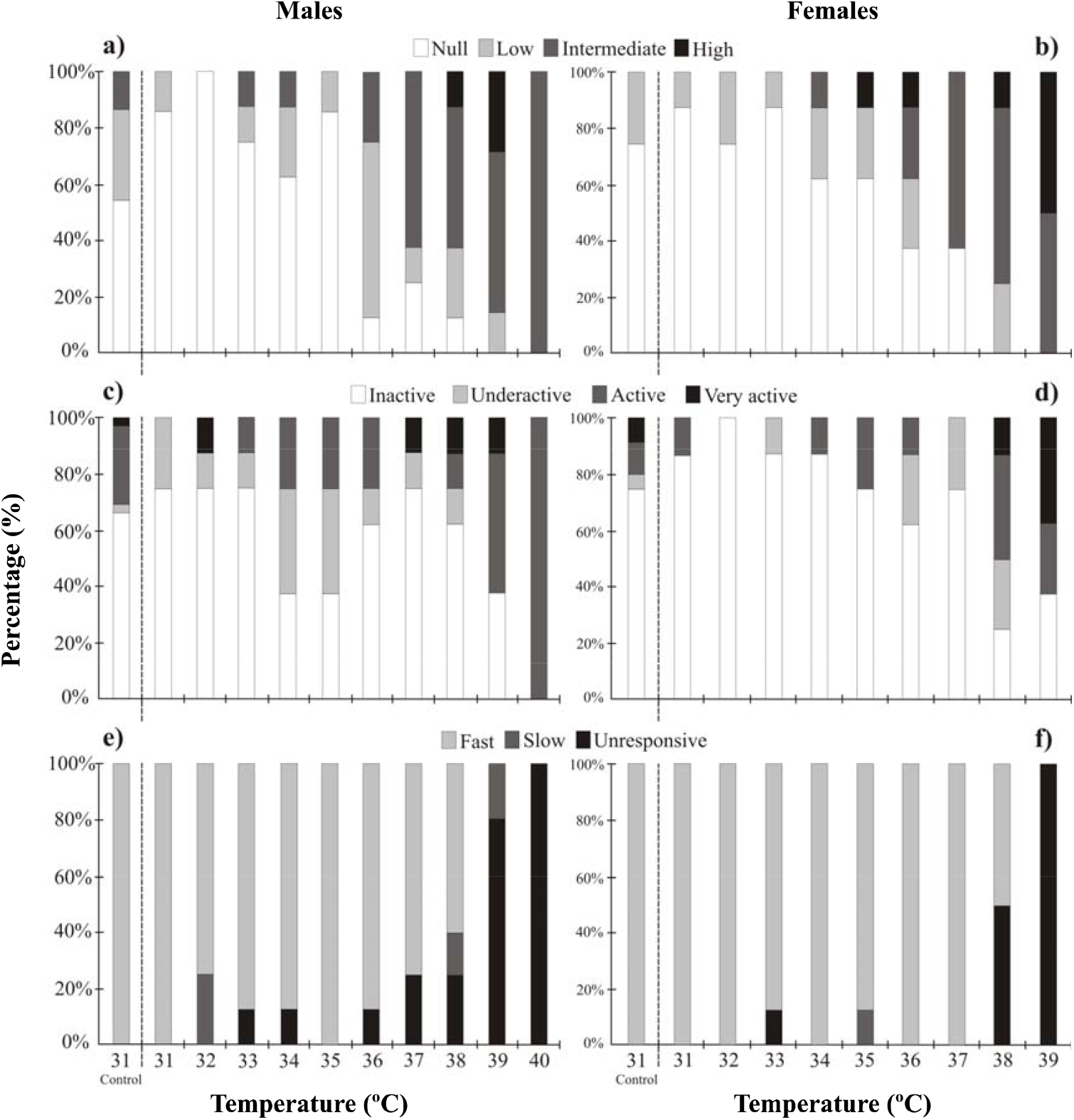
Behavioral responses of 24 *Potamotrygon magdalenae* (12 males and 12 females) individuals to temperature increase, evaluated from: lifting of the disc of (A) males and (B) females; degree of activity of (C) males and (D) females; responses to first stimulus of touch of (E) males and (F) females; *: individual that tolerated up to 40 °C.

Both, in treatments and in control, *P. magdalenae* individuals were mainly inactive, even during most of the temperature increase, up to 37ºC in females (Fig. 2D) and up to 38ºC in males (except at 34ºC and 35ºC, where little activity and inactivity presented the same percentage). Consequently, the stingrays were observed very active, with a constant occurrence beginning at 37ºC in males and an increasing occurrence starting at 38ºC in females.

All control individuals responded immediately to the first stimulus (Fig. 2E,F) and most of the treatment individuals had a rapid response, up to 38°C in males (Fig. 2E) and 37°C in females (Fig. 2F). It was evident that with the increase in temperature the individuals either stopped responding to the first stimulus or did so slowly; this change in response was gradual in males (from 36ºC; Fig. 2E) and abrupt in females (at 38 and 39ºC; Fig. 2F).

### Relationship between CTMax of *Potamotrygon magdalenae* and local and regional temperatures

The highest bottom temperature recorded at Guarinocito Pond was 33,1ºC. This value is 5,9ºC below the CTMax recorded for *P. magdalenae* at this location. Temperatures along Magdalena Basin vary between 12,5ºC and 38,5ºC (mean *±* SD= 27,12 *±* 4,15 ºC). The Guarinocito pond represents the fifth position (among 118 locations) with the highest average temperature (30,1ºC*±* 1,35ºC), distancing 1,5ºC of Malambo swamp, the location with the highest average temperature (SQUALUS Foundation and IDEAM data).

## DISCUSSION

### Critical Thermal Maximum (CTMax) of males and females of *Potamotrygon magdalenae*

The tolerance of fish to low and high temperatures can be affected by multiple factors such as temperature of acclimation, period of acclimation, body size and the rate at which the temperature is changed (Hutchinson, 1976; Lutterschmidt and Hutchison, 1997; Mora and Maya, 2006; Vinagre et al., 2015; Moyano et al., 2017; Desforges et al., 2023). As suggested by Cocking (1959), it can be explained because the time the fish have for acclimatization and the time they were exposed to lethal temperatures, could be affected by the heating rate.

Additionally, fishes acclimated to a high temperature can tolerate higher temperatures than those that are acclimated to a lower temperature and vice versa, which results in the thermal tolerance limit being dependent on the acclimation temperature (Hutchison, 1976; Bennett and Judd, 1992; Mora and Ospina, 2001). In this study the temperature was increased from a single acclimation value and we preferred an increase rate of 1ºC/2h fast enough to induce acute thermal stress responses (Desforges et al., 2023) but slow enough to avoid an inappropriate acclimation due to faster changes in temperature (Mora and Maya, 2006); as Barker et al. (1981) and Hutchinson and Murphy (1985) also suggested, high increase rate can produce long lag between the experimental temperature and the internal temperature of the individual and induce heat-shocks effects, leading to inadequate estimation of CTMax (Lutterschmidt and Hutchinson, 1997).

Furthermore, the use of a dynamic method with a slow increase rate would resulted in the acclimation temperature not having a significant effect on CTMax because fish are able to acclimate to small temperature changes (Elliott and Elliott, 1955; Mora and Ospina, 2001; Mora and Ospina, 2002). The enclosure acclimation period used in this study (24h) was considered sufficient, given that the sampling site and the aquarium water had similar temperatures, which reduce the need for thermal acclimation (Mora and Ospina, 2001). Additionally, our goal was to forecast more realistic outcomes of a sudden heat wave on the local population, therefore we only used field-acclimatized fish.

Although factors as acclimation temperature and acclimation period can influence the tolerance of fish towards high and low temperatures, body size is considered one of the most critical factors (Cox, 1974; Hutchison, 1976; Becker and Genoway, 1979) due to can affect thermal tolerance given by differences in the area/volume ratio or to ontogenetic differences in physiology (Cox, 1974; Hutchison, 1976; Becker and Genoway, 1979; Mora and Ospina, 2004; Mora and Maya, 2006). In the first case, there is lower heat penetration rate in larger organisms, and in the second case, there are differences due to age, sexual maturity and physiological factors related to age (Hutchinson, 1976; Pörtner and Farrell, 2008). Another factor that can potentially affect tolerance limits but has not been widely evaluated is sex (Madeira et al., 2012). Although males and females sizes were similar in this study, sexual dimorphism in size of *P. magdalenae* has been reported, with females reaching larger sizes than males (Lasso et al., 2013). According to this, in the present study there was no relation between CTMax and sex or body size, suggesting that there was no effect of these variables on the tolerance of *P. magdalenae* to high temperatures. From these results, it can be inferred that given a sudden temperature change or a climatic phenomenon that increases temporally the temperature in Guarinocito Pond to values close to the tolerance range of this species (39ºC), *P. magdalenae* individuals could be considerably affected, independently of sex or size. This implies that if both, juveniles and adults are affected, not generational change would be generated in the assessed population (Gotelli, 2001); however, as it has been observed shifts in in thermal performance responses throughout ontogeny (Lear et al., 2019), future studies should include *P. magdalenae* juveniles and neonates.

Temperature records at Guarinocito Pond indicate a bottom maximum temperature (33.1ºC) that is 5.9ºC under the CTMax for the species. This indicates that the stingrays are not being affected by the current temperatures. However, fishermen have reported a worrisome decrease in the water level of the pond, due to a combination of factors that include the temporary closure of the Magdalena River that flowed into the pond, the decrease in precipitation and the increase in environmental temperature (A. Santos, pers. comm.). If those factors that have led to a decrease in water levels continue, it is probable that water temperature could increase enough to go beyond the upper tolerance level of this species, negatively affecting the performance of these animals.

When faced with thermal challenges elasmobranchs move to deep water refugia to reduce stress, i.e. thermotaxis (Di Santo and Bennett, 2011) and to enhance physiological processes throughout small scale movements that could help with survival (Vilmar and Di Santo, 2022). In the study area however, behavioral thermoregulation is not an option for the stingrays, since Guarinocito Pond has homogeneous depths (7,5-9,9 m; Fundación SQUALUS, unpublished data) and the fish cannot ‘escape’ to the river.

Considering the localities in the Magdalena river basin where *P. magdalenae* inhabits and the temperatures recorded in them, it can be inferred that this species has a wide range of temperature (12,5?C-38,5?C) at which individuals can perform their vital functions. Given this wide tolerance range, which varies depending on environmental conditions, it is also necessary to determine if there are one or several populations of this species along the basin, as well as environmental variations among the different localities where the species inhabits.

Although *P. magdalenae* individuals in Guarinocito Pond inhabit relatively below their maximum tolerance limit, considering the data recorded by the IDEAM and SQUALUS Foundation, feeding and growth could decrease significantly at high temperatures, and before the temperatures become lethal (Selong and McMahon, 2001). While crucial traits to elasmobranch vulnerability like locomotor performance can also be affected in different ways in warming scenarios (Vilmar and Di Santo, 2022) Furthermore, other processes involved in the population maintenance (*e.g*., reproduction) could be affected even at temperatures below CTMax, therefore, if the temperature continues to increase, the effect of warming does not necessarily have to be very close to the tolerance limit (Mora and Ospina, 2001). However, it is necessary to differentiate whether increases in temperature occur to short-term (STI) or long-term (LTI), because during STI situations fish need to adapt quickly to altered thermal conditions, while during LTI situations adaptations can be slower. The extent of adaptions to temperature increases depends on genetic variability and characteristics of the life history, such as generational time; however, in a situation of STI or LTI, fish can be seriously threatened with extinction. Animals in LTI have more opportunities for adapting because a higher number of generations can increase genetic variability (Mora and Ospina, 2001). According to this, *P. magdalenae* can be at higher risk when faced with short-term climate changes such as the El Niño phenomenon. However, it is possible that some behavioral modifications can occur to overcome this phenomenon. For example, it has been reported that some bivalves have reproductive strategies focused on decreasing reproductive effort during El Niño, avoiding energy lost in the production of gametes and growth, in order to survive in unfavorable conditions (Urban and Tarazona, 1996).

### Behavior of *Potamotryon magdalenae* individuals against the temperature increase

Lifting of the posterior zone of the disc and/or curling of the fin edges has been recognized as a significant display of stress in freshwater stingrays. It has been reported that if this behavior is prolonged in time then the animal would almost certainly die (Smith et al., 2004). The greater disc lift exhibited at temperatures over 38ºC in males and 35ºC in females, confirms that this behavior is the result of the stress caused by the treatments, being reached faster by females than males. In two thermal tolerance studies conducted on batoids, stress was seen as the tetanic contraction of the disc (muscle spasm manifested as lifting of the disc rear and/or margin of the fins) and was one of the behaviors considered as the sublethal point at which to stop temperature increase, because as mentioned is an indicator of stress (Fangue and Bennett, 2003; Dabruzzi et al., 2012) which supports the selection of this thermal stress response as an end point (Desforges et al., 2023). Additionally, we observed a complete absence of high disc lifting in control individuals, suggesting that this level of disc lifting is indeed related to acute thermal stress.

The increase in temperature causes the elevation of the fish metabolic rate, and thus fish can incur in a mismatch between oxygen demand and supply as suggested by Pörtner and Farrell (2008). This can explain that both males and females showed more activity close to their maximum tolerance limit (higher temperatures), which also could be related with stress ‘restlessness’ in fish (Hill, 2006; Zhang and Kieffer, 2014). Due to aforementioned, individuals in the control could have presented less movement, since the temperature is constant and lower than those of the treatments.

The response of *P. magdalenae* to the touch stimulus was one of the more useful stress indicators in this study. Tactile stimulation of the body, disc or tail has been used in different species of bony fish to trigger an escaping behavior (Beitinger et al., 2000; Mora and Ospina, 2001). The use of this stress indicator may depend on the characteristics of each species; for example, Mora and Ospina (2001) used it for groundfish and verified loss of balance while swimming. It has been observed that when temperature increases the animals cease to respond to stimuli and become more lethargic, losing the ability to escape (burst) from conditions that could be lethal (Beitinger et al., 2000). This was evident in this study, where up to 38 ºC on males and 37 °C on females the response to the first stimulus was usually quick; above this temperature the animals ceased to respond. On the contrary, when animals were not exposed to a temperature increase they responded quickly to the first stimulus, as expected.

## CONCLUSIONS

*Potamotrygon magdalenae* is a species resistant to temperature increase and is currently not affected by temperatures at the study area (Guarinocito Pond). It is probable that if long-term thermal changes occur in the pond, the species could increase its thermal tolerance. However, current environmental changes in the study area have led to a decrease in the water level and an increase in the water temperature that may accelerate, due to lack of water flow from the Magdalena River, the decrease in precipitation and the increase in environmental temperature with short-term climate changes such as El Niño.

The aforementioned, added to the fact that an increase in temperature affects males and females of all development stages equally, can greatly affect the survival of this species in the Guarinocito pond, making it difficult for the population to recover quickly enough after a considerable temperature change.

Of the three variables (disc lifting, activity and the touch response) considered in this study as stress indicators, it is concluded that the last one was a better indicator since the others can be affected by additional factors as water turbidity.

## ACKNOWLEDGMENTS

Authors thank to Andres Felipe Jaramillo for his help in the field phase, Jose Gabriel Perez Rojas and Diego Cordoba for his collaboration with field logistics, Fisherman Association of Guarinocito by obtaining of specimens, IDEAM for provide the data and Squalus Foundation for logistical support and project financing.

## AUTHOR CONTRIBUTIONS

Conceptualization: D.G.M., P.A.M.F.; Methology: D.G.M., E.L.C., P.A.M.F.; Validation: D.G.M., E.L.C., P.A.M.F; Formal analysis: D.G.M., E.L.C., P.A.M.F; Investigation: D.G.M.; Resources: D.G.M., P.A.M.F.; Data Curation: D.G.M.; Writing – original draft preparation: D.G.M.; Writing – review and editing: D.G.M., E.L.C., P.A.M.F, Visualization: D.G.M., E.L.C., P.A.M.F, Supervision: E.L.C., P.A.M.F.; Project administration: D.G.M., P.A.M.F.; Funding acquisition: P.A.M.F.

## DATA AVAILABILITY

The data from the current study are available from the corresponding author on reasonable request.

## LIST OF SYMBOLS AND ABBREVIATIONS

CTMax: Critical Thermal Maximum
CTMin: Critical Thermal Minimum
STI: Short Term Increase
LTI: Long Term Increase
DO: Dissolved Oxygen

